# Stem cell factor acts locally in the bone marrow to promote stem and progenitor cell maintenance

**DOI:** 10.64898/2026.09.28.755094

**Authors:** Maria Angelica Freitas-Cortez, Stefano Comazzetto, Xiang Gao, Bo Shen, Lei Ding, Zhiyu Zhao, Sean J. Morrison

## Abstract

Stem cell factor (SCF) from Leptin Receptor-expressing (LepR^+^) mesenchymal stromal cells and endothelial cells is required for hematopoietic stem and progenitor cell maintenance in bone marrow. Consistent with this, *Scf* is highly expressed by LepR^+^ cells and at lower levels by endothelial cells in bone marrow. However, endothelial cells outside the bone marrow produce circulating SCF, raising the question of whether the SCF that promotes stem and progenitor cell maintenance is produced inside or outside bone marrow. We found that *Scf* deletion in spleen and liver endothelial cells reduces SCF levels in the blood without affecting stem or progenitor numbers in the bone marrow. Moreover, deletion of *Scf* from mesenchymal cells in the long bones, but not vertebrae, of *Prx1*^*Cre*^; *Scf* ^*fl/fl*^ mice depletes stem and progenitor cells in long bones but not vertebrae. SCF thus acts locally within the bones in which it is made to promote stem and progenitor cell maintenance.

**Key Points:**

- *Scf* deletion in endothelial cells outside of BM reduces SCF levels in blood without affecting stem or progenitor cell frequencies in BM
- *Scf* deletion from mesenchymal cells in long bones, but not vertebrae, depletes stem and progenitor cells in long bones but not vertebrae

## Introduction

Adult hematopoiesis is sustained by hematopoietic stem cells (HSCs), multipotent progenitors, and restricted progenitors that reside in the bone marrow. Stem cell factor (SCF, also known as Kit ligand) is critical for the maintenance of HSCs and restricted hematopoietic progenitors^1,2^. *Scf* is highly expressed by perivascular Leptin receptor-expressing (LepR^+^) mesenchymal stromal cells and at lower levels by endothelial cells in the bone marrow^1,3-6^.

HSCs and at least some restricted hematopoietic progenitors reside adjacent to these perivascular LepR^+^ cells^1,3,7^. SCF produced by endothelial cells is selectively required for HSC maintenance^1,3^ while SCF produced by LepR^+^ cells is broadly required for the maintenance of HSCs and restricted hematopoietic progenitors in the bone marrow^3,8^.

SCF is also produced by endothelial cells and other cells outside of the bone marrow^9-11^. This raises the question of whether the SCF that promotes HSC and progenitor maintenance in the bone marrow is produced locally within the bone marrow or outside of the bone marrow. There are membrane-bound and soluble forms of SCF^12^. Membrane-bound SCF may be important for stem and progenitor cell maintenance as HSCs are depleted in *Sl/Sl*^*d*^ mutant mice^13^, which lack membrane-bound SCF but still express soluble SCF^14^; however, the levels of soluble SCF are greatly reduced in these mice^15^ and a recent study found that soluble SCF is more important than membrane bound SCF for HSC maintenance^16^. This emphasizes the need to test if SCF in the blood could have systemic effects on HSC maintenance^16^. Mice with a mixture of wild-type and *Sl/Sl*^*d*^ stromal cells only exhibit normal hematopoiesis in the immediate vicinity of wild-type cells, suggesting that SCF acts locally in the niche^17^. Systemic deletion of membrane-bound, but not soluble, SCF from endothelial cells reduced circulating SCF levels in the blood without having any effect on HSC frequency in the bone marrow^16^. While this observation suggests that HSCs in normal bone marrow do not depend upon SCF from the blood, bones transplanted from fetal *Sl/Sl* mice (which completely lack SCF) into wild-type mice had normal HSC frequency and reduced erythroid progenitor frequency in the marrow of the transplanted bones three weeks after transplantation^16^. This suggests HSCs can be sustained, at least transiently, by soluble SCF produced outside the bone marrow when no SCF is made inside the bone marrow. It is unclear if SCF from the blood contributes to HSC maintenance in the bone marrow under physiological conditions when SCF is synthesized in the bone marrow.

## Study Design

To address this question, we deleted *Scf* from endothelial cells in the spleen and liver using *Lyve1*^*Cre*^ (ref^18^) or *Vav1*^*Cre*^ (ref^19^), then examined the effects on SCF levels in the blood and bone marrow as well as HSC and progenitor numbers in the bone marrow. We also deleted *Scf* from mesenchymal cells using *Prx1*^*Cre*^ (ref^20^), which recombines in the long bones but not in vertebrae, then examined the effects on SCF levels in the blood, long bones, and vertebrae as well as HSC and progenitor numbers in the long bones, and vertebrae. Detailed methods are provided in the supplemental Methods.

## Results and Discussion

The *Vav1*^*Cre*^ allele we used recombines in endothelial cells in the spleen and liver but not in endothelial cells in bone marrow^9^. Spleen and liver endothelial cells express *Scf* ^*9,22*^. Vav1-cre also recombines in hematopoietic cells, which express little or no *Scf* ^1^. Prior studies found that *Vav1*^*Cre*^; *Scf* ^*fl/fl*^ and littermate control mice did not significantly differ in terms of HSC or restricted progenitor numbers in the bone marrow^1,9^, suggesting that neither hematopoietic cells nor endothelial cells outside of the bone marrow were functionally important sources of SCF for HSC or progenitor maintenance in the bone marrow. We independently tested this, first confirming by qRT-PCR that *Scf* transcript levels were significantly reduced in endothelial cells from the spleen, but not the bone marrow, of *Vav1*^*Cre*^; *Scf* ^*fl/fl*^ as compared to *Scf* ^*fl/fl*^ controls (Supplemental Figure 2A and 2B)^9^. By ELISA analysis, SCF levels were 40% lower in blood serum from *Vav1*^*Cre*^; *Scf* ^*fl/fl*^ mice as compared to *Scf* ^*fl/fl*^ controls (Figure 1A), but SCF levels in bone marrow fluid from femurs and tibias did not significantly differ between *Vav1*^*Cre*^; *Scf* ^*fl/fl*^ and control mice (Figure 1B). Even modest reductions in SCF levels significantly reduce HSC and progenitor numbers in the bone marrow^1,23^. However, bone marrow cells from *Vav1*^*Cre*^; *Scf* ^*fl/™*^ and littermate control mice gave similar levels of donor cell reconstitution upon competitive transplantation into irradiated recipients (Supplemental Figure 2C). These observations, and published data^1,9^, suggest that HSCs and progenitors in the bone marrow do not depend upon SCF from the blood.

**Figure 1.**
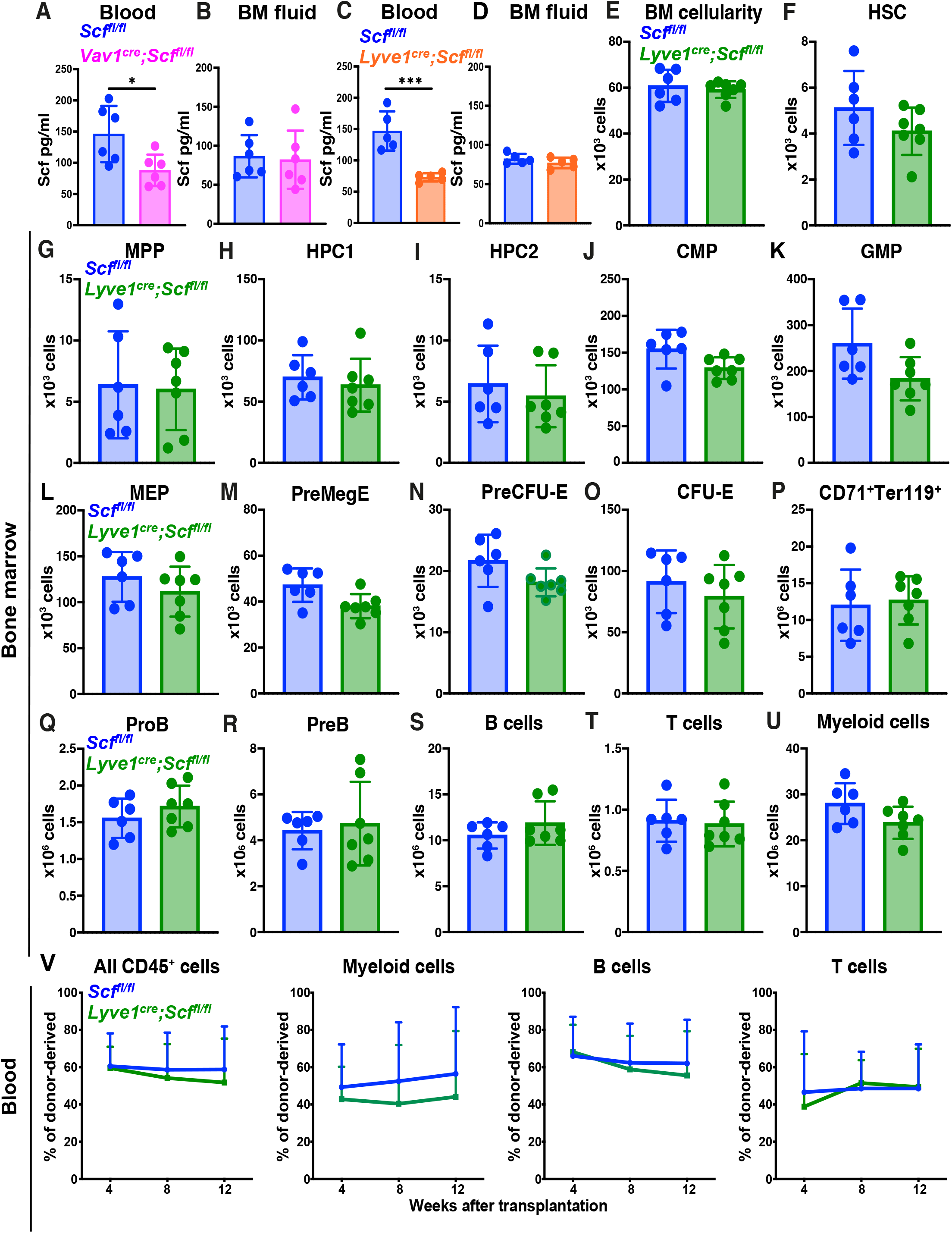
Deletion of *Scf* from spleen and liver endothelial cells reduces SCF levels in the blood without affecting the frequencies of hematopoietic stem or progenitor cells in the bone marrow. (A, B) SCF levels in blood serum (A) and bone marrow (BM) fluid collected from the femurs and tibias (B) of *Vav1*^*Cre*^; *Scf* ^*fl/fl*^ and *Scf* ^*fl/fl*^ control mice (a total of 6 mice per genotype from 6 independent experiments). (C, D) SCF levels in blood plasma (C) and BM fluid (D) collected from the femurs and tibias of *Lyve1*^*Cre*^; *Scf* ^*fl/fl*^ and *Scf* ^*fl/fl*^ control mice (a total of 5 mice per genotype from 3 independent experiments). (E-U) Bone marrow cellularity (E), as well as the numbers of HSCs (F), MPPs (G), HPC1 (H), HPC2 (I), CMPs (J), GMPs (K), MEPs (L), PreMegEs (M), PreCFU-Es (N), CFU-Es (O), CD71^+^Ter119^+^ erythroid cells (P), ProB cells (Q), PreB cells (R), B cells (S), T cells (T), and myeloid cells (U) in bone marrow from two femurs and two tibias of *Lyve1*^*Cre*^; *Scf* ^*fl/fl*^ and *Scf* ^*fl/fl*^ control mice (a total of 6-7 mice per genotype from 4 independent experiments). (V) Donor cell reconstitution in the blood of irradiated mice after transplantation of 500,000 donor bone marrow cells from *Lyve1*^*Cre*^; *Scf* ^*fl/fl*^ or *Scf* ^*fl/fl*^ control mice along with 500,000 competing wild-type cells (a total of 14-15 recipient mice per genotype transplanted with cells from 3 donors per genotype in 3 independent experiments). All data represent mean ± standard deviation. Each dot in panels A-U represents a different mouse. Statistical significance was assessed using a matched samples two-way ANOVA followed by Sidak’s multiple comparisons adjustment (A, B), Student’s *t*-tests followed by Holm-Sidak’s multiple comparisons adjustments (C, D, and F-U), a Student’s *t*-test (E), or non-parametric analysis of longitudinal data (nparLD) with the False Discovery Rate (FDR) method for multiple comparisons adjustments to test differences in overall reconstitution (V). All statistical tests were 2-sided (*, P<0.05; ***, P<0.001).

We independently assessed this using *Lyve1*^*Cre*^, which recombines in fetal endothelial cells^24^ and lymphatic endothelial cells^18^. In 3-month-old *Lyve1*^*Cre*^; *tdTomato* reporter mice we observed recombination (Tomato^+^) in 11 ± 4.8% of bone marrow endothelial cells, 83 ± 4.0% of spleen endothelial cells, and 88 ± 1.4% of liver endothelial cells (Supplemental Figure 3A). Consistent with this, *Scf* transcript levels were significantly reduced in endothelial cells from the spleen, but not bone marrow, of adult *Lyve1*^*Cre*^;*Scf* ^*fl/fl*^ mice as compared to *Scf* ^*fl/fl*^ controls (Supplemental Figure 3B and 3C). Three month-old *Lyve1*^*Cre*^; *Scf* ^*fl/fl*^ mice exhibited approximately 50% lower SCF levels in blood plasma (Figure 1C), but not bone marrow fluid, relative to *Scf* ^*fl/fl*^ controls (Figure 1D).

Three month-old *Lyve1*^*Cre*^; *Scf* ^*fl/fl*^ mice and *Scf* ^*fl/fl*^ controls did not significantly differ in terms of blood cell counts (Supplemental Figure 3D-O), spleen cellularity (Supplemental Figure 3P), spleen HSC and MPP numbers (Supplemental Figure 3Q and 3R), bone marrow cellularity, or the numbers of HSCs, MPPs, restricted hematopoietic progenitors, or myeloid, B, or T cells in the bone marrow (Figure 1E-U). Moreover, bone marrow cells from *Lyve1*^*Cre*^; *Scf* ^*fl/fl*^ mice and *Scf* ^*fl/fl*^ controls gave similar levels of multilineage reconstitution upon competitive transplantation into irradiated mice (Figure 1V). This further suggests that HSCs and progenitors in normal bone marrow do not depend upon SCF from the blood.

To test whether SCF produced in the bone marrow acts locally or systemically to promote stem and progenitor cell maintenance, we generated *Prx1*^*Cre*^; *Scf* ^*fl/fl*^ mice. *Prx1*^*Cre*^ recombines in mesenchymal cells during fetal development, including in the cells that give rise to LepR^+^ bone marrow cells, in long bones (femurs and tibias) but not in vertebrae or other parts of the axial skeleton^20^. Consistent with this, *Scf* transcript levels were reduced in LepR^+^ bone marrow cells from the long bones, but not vertebrae, of *Prx1*^*Cre*^;*Scf* ^*fl/fl*^ as compared to *Scf* ^*fl/fl*^ controls (Supplemental Figure 4A and 4B). By ELISA analysis, *Prx1*^*Cre*^; *Scf* ^*fl/fl*^ mice exhibited reduced SCF levels in femur and tibia bone marrow fluid as compared to *Scf* ^*fl/fl*^ controls (approximately 40% lower; Figure 2A) but not in vertebral bone marrow fluid (Figure 2B) or blood (Figure 2C). This indicates that SCF levels in the bone marrow are primarily regulated locally.

**Figure 2.**
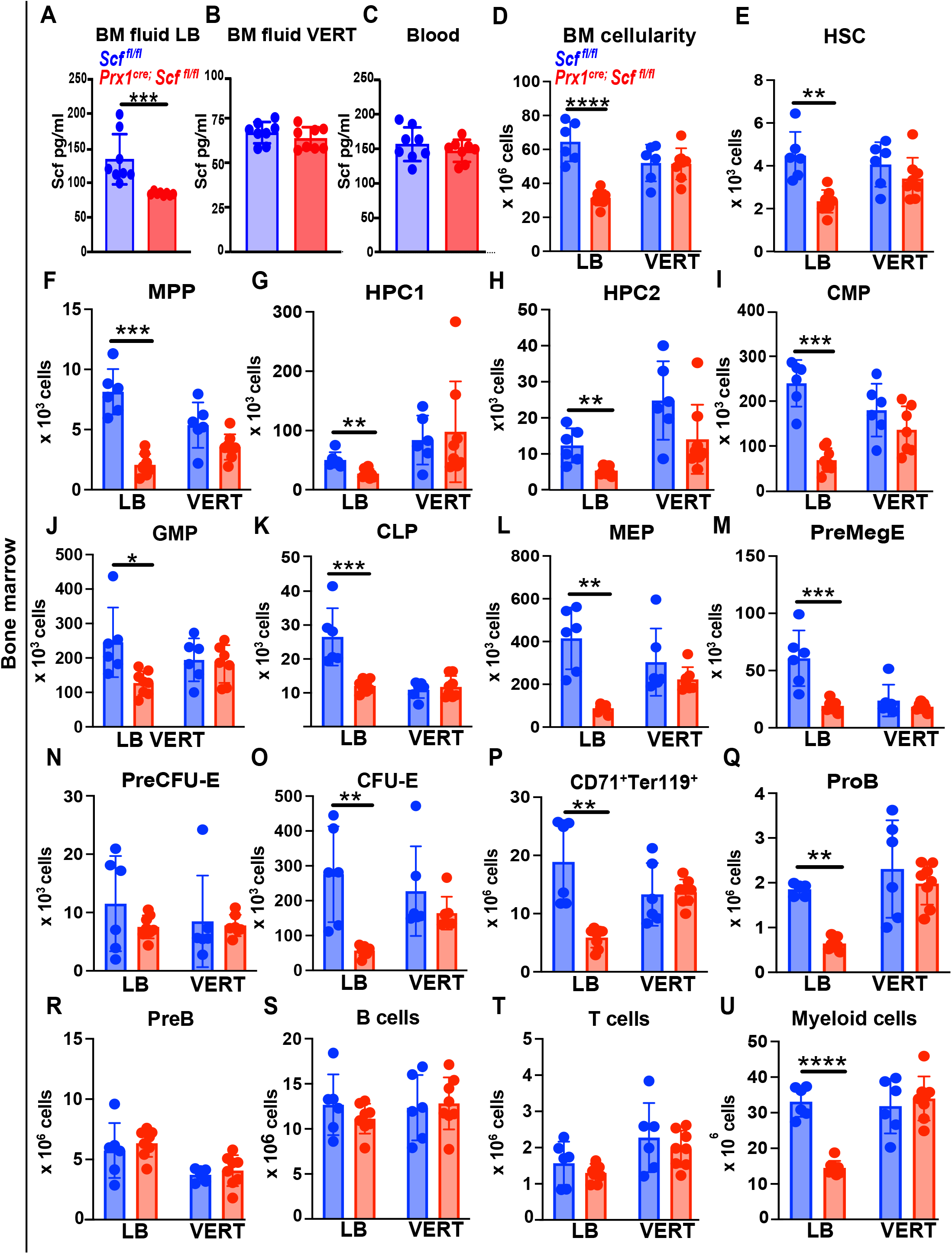
SCF acts locally within the bone marrow compartment in which it is produced to promote the maintenance of HSCs and restricted hematopoietic progenitors. *Prx1*^*Cre*^ recombines in mesenchymal cells, including LepR^+^ cells, in femurs and tibias but not in vertebrae^20^. (A-C) SCF protein levels in bone marrow fluid from the long bones (LB; femurs and tibias) (A) or vertebrae (VERT) (B), as well as from blood plasma (C) from *Prx1*^*Cre*^; *Scf* ^*fl/fl*^ and *Scf* ^*fl/fl*^ control mice (a total of 8 mice per genotype from 2 independent experiments). (D-U) Bone marrow cellularity (D) and the numbers of HSCs (E), MPPs (F), HPC1 (G), HPC2 (H), CMPs (I), GMPs (J), CLPs (K), MEPs (L), PreMegEs (M), PreCFU-Es (N), CFU-Es (O), CD71^+^Ter119^+^ erythroid cells (P), ProB cells (Q), PreB cells (R), B cells (S), T cells (T), and myeloid cells (U) in bone marrow from the long-bones and vertebrae of *Prx1*^*Cre*^; *Scf* ^*fl/fl*^ and control *Scf* ^*fl/fl*^ mice (a total of 6-8 mice per genotype from 2 independent experiments). Each dot represents a different mouse, and all data represent mean ± standard deviation. Statistical significance was assessed using Student’s *t*-tests or Mann-Whitney tests followed by Holm-Sidak’s multiple comparisons adjustments (A-C, D, and E-U). All statistical tests were 2-sided (*, P <0.05; **, P<0.01; ***, P<0.001; ****, P<0.0001).

In vertebral bone marrow, *Prx1*^*Cre*^; *Scf* ^*fl/fl*^ mice and *Scf* ^*fl/fl*^ controls did not significantly differ in terms of bone marrow cellularity, or the numbers of HSCs, MPPs, restricted hematopoietic progenitors, or myeloid, B, or T cells (Figure 2D-U). Furthermore, vertebral bone marrow cells from *Prx1*^*Cre*^; *Scf* ^*fl/fl*^ mice and *Scf* ^*fl/fl*^ littermate controls did not significantly differ in their capacity to give long-term multilineage reconstitution upon competitive transplantation into irradiated mice (Figure 3A). Conversely, in femurs and tibias, *Prx1*^*Cre*^; *Scf* ^*fl/fl*^ mice had significantly reduced numbers of bone marrow cells, HSCs, MPPs, and most restricted progenitors and myeloid cells as compared to *Scf* ^*fl/fl*^ controls (Figure 2D-U). Femur and tibia bone marrow cells from *Prx1*^*Cre*^; *Scf* ^*fl/fl*^ mice also gave significantly lower levels of myeloid, B, and T cell reconstitution upon competitive transplantation into irradiated mice (Figure 3B). Therefore, SCF produced by bone marrow mesenchymal cells regulates stem and progenitor cell maintenance by acting locally in the bone marrow compartment in which it is made.

**Figure 3.**
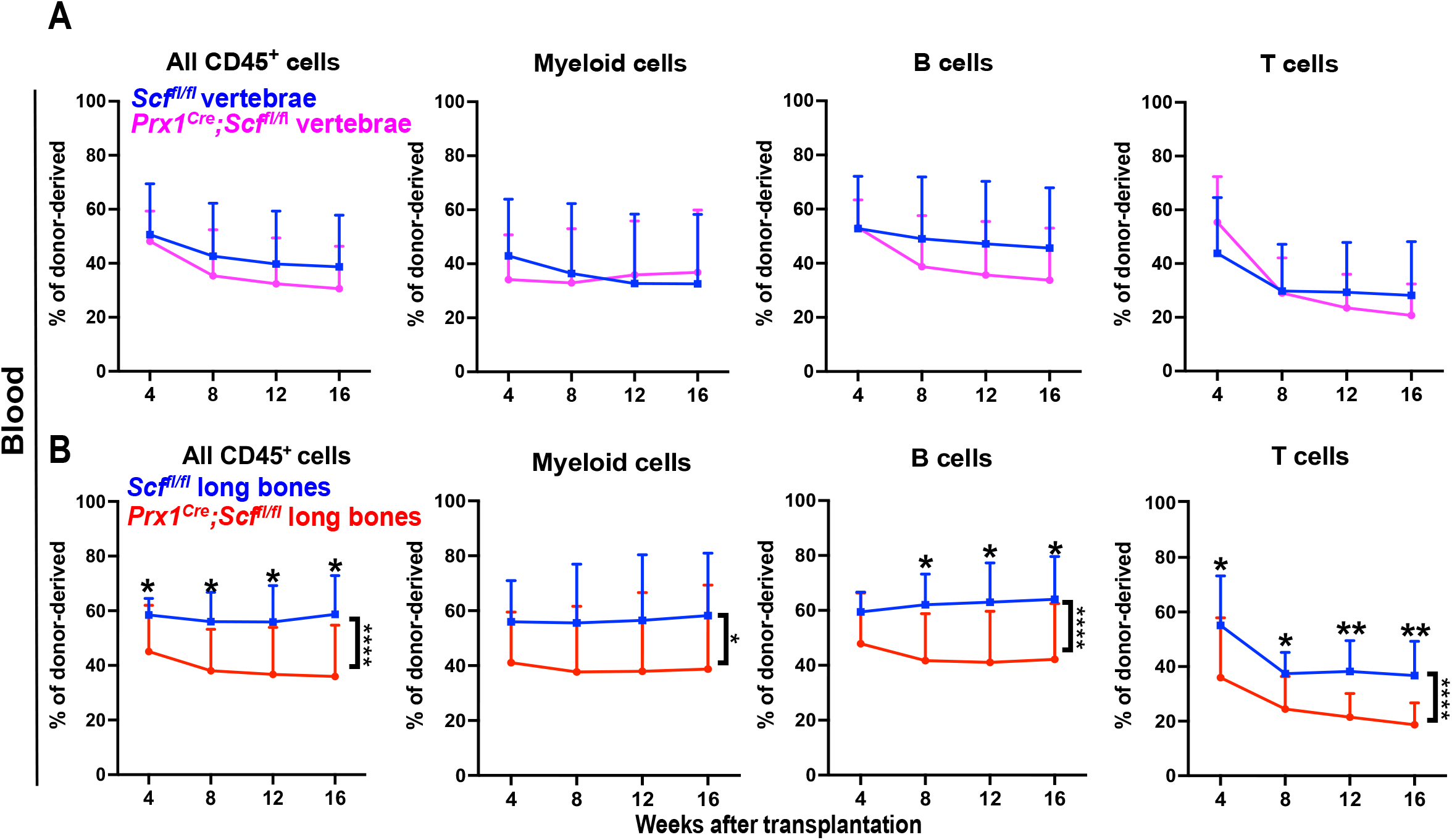
*Prx1*^*Cre*^; *Scf* ^*fl/fl*^ mice have reduced reconstituting activity in bone marrow from femurs and tibias, but not vertebrae, upon competitive transplantation into irradiated mice. (A-B) Donor-derived reconstitution of CD45^+^ cells, myeloid, B, and T cells in the blood of recipient mice after competitive transplantation of 500,000 donor bone marrow cells from the vertebrae (A) or femurs and tibias (long bones; B) of *Prx1*^*Cre*^; *Scf* ^*fl/fl*^ or *Scf* ^*fl/fl*^ control mice, along with 500,000 competing wild-type cells, into irradiated recipients (a total of 8-10 recipient mice per genotype transplanted with cells from 2-3 donors per genotype in 2 experiments). All data represent mean ± standard deviation. Statistical significance was assessed using Student’s *t*-tests or Mann-Whitney tests followed by Holm-Sidak’s multiple comparisons adjustments for the time-point differences or nparLD tests followed by FDR multiple comparisons adjustments to test differences in overall reconstitution (A, B). All statistical tests were 2-sided (*, P <0.05; **, P<0.01; ****, P<0.0001).

Deletion of *Scf* from spleen and liver endothelial cells reduced SCF levels in the blood without affecting stem or progenitor cell numbers in bone marrow (Figure 1), though it is possible that Vav1-cre or Lyve1-cre also recombined in additional *Scf*-expressing cells outside the bone marrow. Either way, the observation that SCF can be depleted from the blood without affecting HSC or progenitor numbers in the bone marrow suggests that stem and progenitor cells in normal bone marrow depend mainly on SCF synthesized in the bone marrow. However, if SCF is completely eliminated from the bone marrow, SCF from the blood may at least transiently rescue HSC maintenance^16^.

Our data do not argue against the conclusion that soluble SCF is important for stem and progenitor cell maintenance^16^. Indeed, the companion study by Zhang et al. found that SCF produced within dorsal vertebrae can promote the maintenance of HSCs and progenitors in ventral vertebrae, implicating soluble SCF^25^. Nonetheless, both of our studies suggest that, under normal circumstances, SCF produced by mesenchymal stromal cells within the bone marrow promotes HSC and restricted progenitor maintenance by acting locally within the bone marrow compartment in which it is produced.

## Data sharing statement

For original data, contact

## Data and code availability

All data reported in this paper will be shared by the lead contact upon reasonable request.

## Materials availability

A material transfer agreement is required to obtain mice in accordance with the regulations of the University of Texas Southwestern Medical Center.

## Acknowledgments

S.J.M. is a Howard Hughes Medical Institute Investigator, the Mary McDermott Cook Chair in Pediatric Genetics, the Kathryn and Gene Bishop Distinguished Chair in Pediatric Research, the director of the Hamon Laboratory for Stem Cells and Cancer, and a Cancer Prevention and Research Institute of Texas Scholar. This work was supported by the National Institutes of Health (DK118745) and the Kleberg Foundation. We thank M. Ortiz and the Moody Foundation Flow Cytometry Facility, Daniel Cassidy for mouse colony management and the BioHPC High-Performance Facility Cloud at UT Southwestern Medical Center for providing computational resources. This article is subject to HHMI’s Open Access to Publications policy. HHMI lab heads have previously granted a nonexclusive CC BY 4.0 license to the public and a sublicensable license to HHMI in their research articles. Pursuant to those licenses, the author-accepted manuscript of this article can be made freely available under a CC BY 4.0 license immediately upon publication.

## Authorship Contributions

M.A.F.C., S.C. and S.J.M. conceived the project, designed and interpreted experiments. M.A.F.C., S.C., X.G., B.S., and L.D. performed all experiments. Z.Z. performed statistical analyses. M.A.F.C., S.C. and S.J.M. wrote the manuscript.

## Supplemental Material

**Supplemental Figure 1. Markers and flow cytometry gates used to identify bone marrow hematopoietic stem and progenitor cell populations**. (A) Representative flow cytometry gates used to identify hematopoietic stem and progenitor cell populations in the bone marrow, including LSK cells, LKS-myeloid progenitors, HSCs, multipotent progenitor cells (MPPs), hematopoietic progenitor cells 1 (HPC1), HPC2, common myeloid progenitors (CMPs), granulocyte-monocyte progenitors (GMPs), megakaryocyte-erythroid progenitors (MEPs), colony-forming units-erythroid (CFU-E), PreCFU-E, and PreMegE cells. (B) Representative flow cytometry gates used to identify differentiating hematopoietic cell populations in the bone marrow, including B cells, ProB cells, PreB cells, T cells, CD71^+^Ter119^+^ erythroid cells, and myeloid cells. (C) Representative flow cytometry gates used to identify common lymphoid progenitors (CLP). The HSC, MPP, HPC1, and HPC2 populations were identified as described by Oguro et al.^21^, whereas CLP, CMP, GMP, MEP, PreMegE, PreCFU-E, and CFU-E populations were identified as described by Comazzetto et al^3^ and Pronk et al^25^.

**Supplemental Figure 2. *Vav1-Cre*-mediated *Scf* deletion reduces *Scf* transcript levels in spleen, but not bone marrow, endothelial cells and does not significantly affect the multilineage reconstituting potential of bone marrow cells upon transplantation into irradiated mice**. (A, B) Relative *Scf* transcript levels in spleen (A) and bone marrow (B) endothelial cells from *Vav1*^*Cre*^; *Scf* ^*fl/fl*^ and *Scf* ^*fl/fl*^ control mice (a total of 3 mice per genotype from 3 independent experiments; each dot represents a different mouse). (C) Donor-derived reconstitution of CD45^+^ cells, myeloid, B, and T cells in the blood of recipient mice after competitive transplantation of 500,000 donor bone marrow cells from the femurs and tibias of *Vav1*^*Cre*^; *Scf*^*fl/-*^ mice, *Scf*^*+/+*^ mice, and *Scf*^*+/-*^ mice, along with 500,000 competing wild-type cells, into irradiated recipients (5 recipient mice per genotype). Statistical significance was assessed using one-sample *t*-tests followed by Holm-Sidak multiple comparisons adjustments (A, B) or nparLD followed by FDR multiple comparisons adjustments (C). All data represent mean ± standard deviation and statistical tests were 2-sided (*, P < 0.05; **, P < 0.01; ***, P < 0.001).

**Supplemental Figure 3. *Lyve1-Cre* recombines in spleen and liver, but not bone marrow, endothelial cells and *Lyve1***^***Cre***^**; *Scf*** ^***fl/fl***^ **mice have normal steady-state hematopoiesis**. (A) The percentage of VE-cadherin^+^ endothelial cells that were tdTomato+ in bone marrow, spleen, and liver of 3-month-old *Lyve1*^*Cre*^; *tdTomato* mice (a total of 5–6 mice from four independent experiments). (B, C) Relative *Scf* transcript levels in spleen (B) and bone marrow (C) endothelial cells from *Lyve1*^*Cre*^; *Scf* ^*fl/fl*^ and *Scf* ^*fl/fl*^ control mice (a total of 3 mice per genotype from 3 independent experiments). (D-O) White blood cell (D) and red blood cell (E) counts, hemoglobin levels (F) and hematocrit (G), mean corpuscular volume (H), mean corpuscular hemoglobin (I), mean corpuscular hemoglobin concentration (J), red cell distribution width (K), and neutrophil (L), lymphocyte (M), monocyte (N) and platelet (O) counts in the blood of *Lyve1*^*Cre*^; *Scf* ^*fl/fl*^ and littermate control (*Scf* ^*fl/fl*^) mice. (P-R) Spleen cellularity (P) as well as the numbers of HSCs (Q) and MPPs (R) in the spleen of *Lyve1*^*Cre*^; *Scf* ^*fl/fl*^ and littermate control mice. Data are from a total of 6-7 mice from 4 independent experiments). All data represent mean ± standard deviation and each dot represents a different mouse. Statistical significance was assessed using a linear mixed-effects model followed by Dunnett’s multiple comparisons adjustment (A) one-sample *t*-tests followed by Holm-Sidak’s multiple comparisons adjustment (B, C) Student’s *t*-tests or Mann-Whitney tests followed by Holm-Sidak’s multiple comparisons adjustments (D-O, and Q-R) or a Student’s *t*-test (P). All statistical tests were 2-sided.

**Supplemental Figure 4. *Prx1-Cre*-mediated recombination of *Scf* reduces *Scf* transcript levels in LepR**^+^ **bone marrow stromal cells in long bones (femurs and tibia) but not vertebrae**. (A, B) Relative *Scf* transcript levels in LepR^+^ bone marrow stromal cells from long bones (A) versus vertebrae (B) of *Prx1*^*Cre*^; *Scf* ^*fl/fl*^ and *Scf* ^*fl/fl*^ control mice (a total of 3 mice per genotype from 3 independent experiments). All data represent mean ± standard deviation and each dot represents a different mouse. Statistical significance was assessed using one-sample *t*-tests followed by Holm-Sidak’s multiple comparisons adjustment (A, B).

## Supplemental Methods

### Mice

The *Scf* ^*fl*^ allele (RRID:IMSR_JAX:017861) was generated in our laboratory^1^. *Lyve1*^*Cre*^ (RRID:IMSR_JAX:012601)^18^, *Vav1*^*Cre*^ (RRID:IMSR_JAX:008610)^19^, *Prx1*^*Cre*^ (RRID:IMSR_JAX:005584)^20^, and *Rosa26LSL-tdTomato* reporter mice (B6.Cg-Gt(ROSA)26Sortm14(CAG-tdTomato)Hze/J; RRID:IMSR_JAX:007914) were obtained from Jackson Laboratory. All mice were maintained on a C57BL/Ka background. We used littermate or age-matched mice of both sexes that were 8-12 weeks or 4-6 months old in experiments. Mice were housed in AAALAC-accredited, specific-pathogen-free animal care facilities at UT Southwestern Medical Center (UTSW). All procedures were approved by the UTSW Institutional Animal Care and Use Committee.

### Flow cytometric analysis of hematopoietic cells and transplantation assays

Bone marrow cells were obtained from femurs and tibias and analyzed by flow cytometry as previously described^3^. Vertebral bone marrow was obtained by removing the spinal cord and associated soft tissue, crushing vertebral bones in staining medium (Ca^2+^/Mg^2+^-free Hanks’ balanced salt solution (HBSS) supplemented with 2% heat-inactivated bovine serum), and filtering to generate single-cell suspensions. Antibodies and gating strategies for analysis of hematopoietic stem and progenitor cells, as well as mature hematopoietic cells, were as described previously^3^. Dead cells were excluded by propidium iodide staining. Cells were analyzed using a BD FACSLyric cytometer. For competitive transplantation, 500,000 unfractionated bone marrow cells from donor (CD45.2) and competitor (CD45.1) mice were mixed and injected into the retro-orbital venous sinus of irradiated (CD45.1/CD45.2) recipients. Peripheral blood chimerism was assessed every 4 weeks by flow cytometry as previously described^3,24,25^. HSC, MPP, HPC1, and HPC2 populations were identified as previously described by Oguro et al.^21^, whereas CLP, CMP, GMP, MEP, PreMegE, PreCFU-E, and CFU-E populations were identified as described by Comazzetto et al^3^ and Pronk et al^25^. Markers and flow cytometry gates used to identify bone marrow hematopoietic stem and progenitor cell populations are shown in Supplemental Table 1 and Supplemental Figure 1.

### ELISA analysis

Blood was collected into ethylenediaminetetraacetic acid (EDTA)-containing tubes (to prevent clotting) to obtain plasma or empty tubes to obtain serum. The samples were centrifuged at 2,400 x g for 5 minutes to separate plasma or serum from the cellular fraction or clot, respectively. SCF concentrations were measured in the plasma or serum using the Mouse SCF Quantikine ELISA Kit (R&D Systems, MCK00) according to the manufacturer’s instructions. SCF levels were measured in blood serum, blood plasma, or bone marrow fluid from long bones (femurs and tibias) or vertebrae. Bone marrow fluid was obtained from long bones by cutting the ends off femurs and tibias, then centrifuging the bones at 2,400 x g for 5 minutes to centrifuge the bone marrow out of the bones and into 10 ul of phosphate-buffered saline (PBS), then the supernatant was collected. Vertebral bone marrow fluid was prepared by crushing vertebrae in PBS, followed by centrifugation to collect the supernatant.

### Isolation of endothelial cells and LepR^+^ stromal cells and RT-qPCR analysis

Bone marrow and spleen endothelial cells and bone marrow LepR^+^ stromal cells were isolated as previously described^2,9^. Bone marrow cells were obtained from crushed femurs and tibias and enzymatically dissociated in HBSS containing DNase I (200 U/mL), Dispase (4 mg/mL), and collagenase I (3 mg/mL) for 30 minutes at 37°C. Spleens were mechanically dissociated and enzymatically digested as previously described^2,9^. Cells were stained with biotinylated anti-LepR (AF497; R&D Systems), anti-CD45 (30-F11; APC-eFluor 780; eBioscience), anti-Ter119 (TER-119; APC-eFluor 780; eBioscience), anti-CD31 (clone 390; PE/Cyanine7-conjugated, BioLegend, 102418), and streptavidin (BV421-conjugated, 405226; BioLegend). Dead cells were excluded using Ghost Dye Red 780 (13-0865-T100; Tonbo Biosciences). Cells were sorted using a FACSAria II (BD Biosciences) directly into TRIzol. Total RNA was extracted according to the manufacturer’s instructions, and cDNA was synthesized using the iScript cDNA Synthesis Kit (Bio-Rad). Quantitative real-time PCR was performed using iTaq Universal SYBR Green Supermix (Bio-Rad). *Scf* expression was normalized to *Actb* (which encodes β-actin), and relative expression was calculated using the 2^?ΔΔCt^ method. Primers were as previously described by Ding et al.^1^: *Scf*, forward 5′-TTGTTACCTTCGCACAGTGG-3′ and reverse 5′-AATTCAGTGCAGGGTTCACA-3′; *Actb*, forward 5′-GCTCTTTTCCAGCCTTCCTT-3′ and reverse 5′-CTTCTGCATCCTGTCAGCAA-3′.

### Statistical methods

In each type of experiment, multiple mice were tested in multiple independent experiments performed on different days. Mice were allocated to experiments randomly and samples processed in an arbitrary order, but formal randomization techniques were not used. No formal blinding was applied when performing the experiments or analyzing the data. Sample sizes were not pre-determined based on statistical power calculations but were based on our experience with these assays. No data were excluded. Prior to analyzing the statistical significance of differences among treatments, we tested whether data were normally distributed and whether variance was similar among groups.

To test for normality, we performed Shapiro–Wilk tests when 3≤n<20 or D’Agostino-Pearson omnibus normality tests when n≥20. To test whether variability significantly differed among groups we performed *F*-tests (for experiments with two groups) or Levene’s median tests (for experiments with more than two groups). When the data significantly deviated from normality or variability significantly differed among groups, we log2-transformed the data and tested again for normality and variability. If the transformed data no longer significantly deviated from normality and equal variability, we performed parametric tests on the transformed data. If log2-transformation was not possible or the transformed data still significantly deviated from normality or equal variability, we performed non-parametric tests on the non-transformed data.

When data or log2-transformed data were normal and equally variable, statistical analysis was performed using Student’s *t*-tests (when there were two groups), matched samples two-way ANOVAs (when there were two or more groups), or mixed effects analyses (when there were missing values but the data otherwise met the assumptions for matched samples ANOVAs). When the data and log2-transformed data were abnormal or unequally variable, statistical analysis was performed using Mann-Whitney tests (when there were two groups), Kruskal-Wallis tests (when there were more than two groups), or nparLD tests (when there were two or more groups measured at multiple time points). P-values from multiple comparisons were adjusted using Tukey’s (for all possible comparisons), Sidak’s (for planned comparisons), or Dunnett’s methods (for comparisons between the control and treatment groups) following ANOVAs or mixed effects analyses, Dunn’s method following Kruskal-Wallis tests, or Benjamini-Hochberg’s False Discovery Rate (FDR) method following nparLD tests. Holm-Sidak’s method was used to adjust comparisons following *t*-tests or Mann-Whitney tests. All statistical tests were two-sided where applicable. All data represent mean ± standard deviation. Statistical tests were performed using GraphPad Prism V11.0.1 or R 4.5.0.

