## Supplementary Material for "Stem cell factor acts locally in the bone marrow to promote stem and progenitor cell maintenance"

**SUPPLEMENTAL MATERIAL**
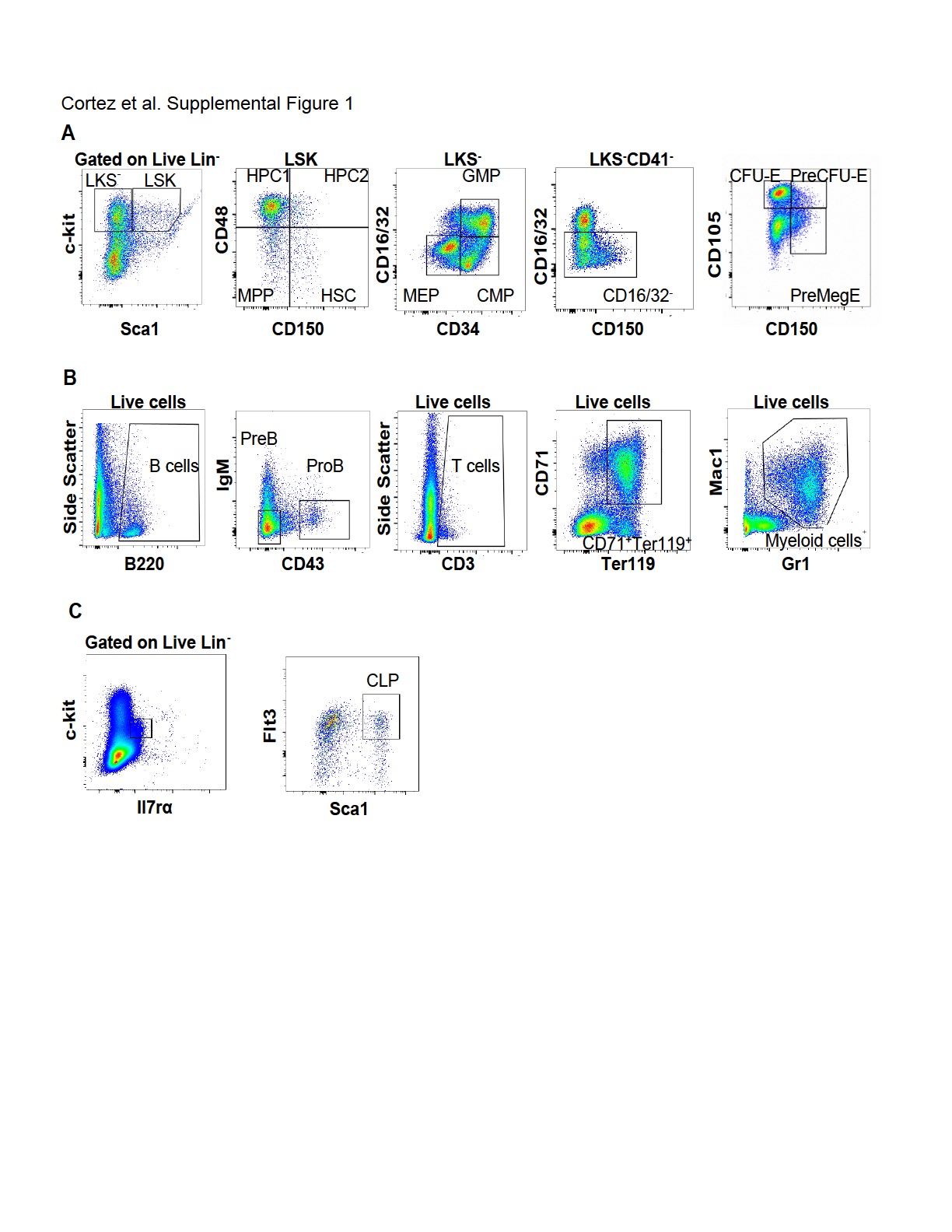

**Supplemental Figure 1. Markers and flow cytometry gates used to identify bone marrow hematopoietic stem and progenitor cell populations.** (A) Representative flow cytometry gates used to identify hematopoietic stem and progenitor cell populations in the bone marrow, including LSK cells, LKS- myeloid progenitors, HSCs, multipotent progenitor cells (MPPs), hematopoietic progenitor cells 1 (HPC1), HPC2, common myeloid progenitors (CMPs), granulocyte-monocyte progenitors (GMPs), megakaryocyte-erythroid progenitors (MEPs), colony-forming units-erythroid (CFU-E), PreCFU-E, and PreMegE cells. (B) Representative flow cytometry gates used to identify differentiating hematopoietic cell populations in the bone marrow, including B cells, ProB cells, PreB cells, T cells, CD71^+^Ter119^+^ erythroid cells, and myeloid cells. (C) Representative flow cytometry gates used to identify common lymphoid progenitors (CLP). The HSC, MPP, HPC1, and HPC2 populations were identified as described by Oguro et al.^21^, whereas CLP, CMP, GMP, MEP, PreMegE, PreCFU-E, and CFU-E populations were identified as described by Comazzetto et al^3^ and Pronk et al^26^.

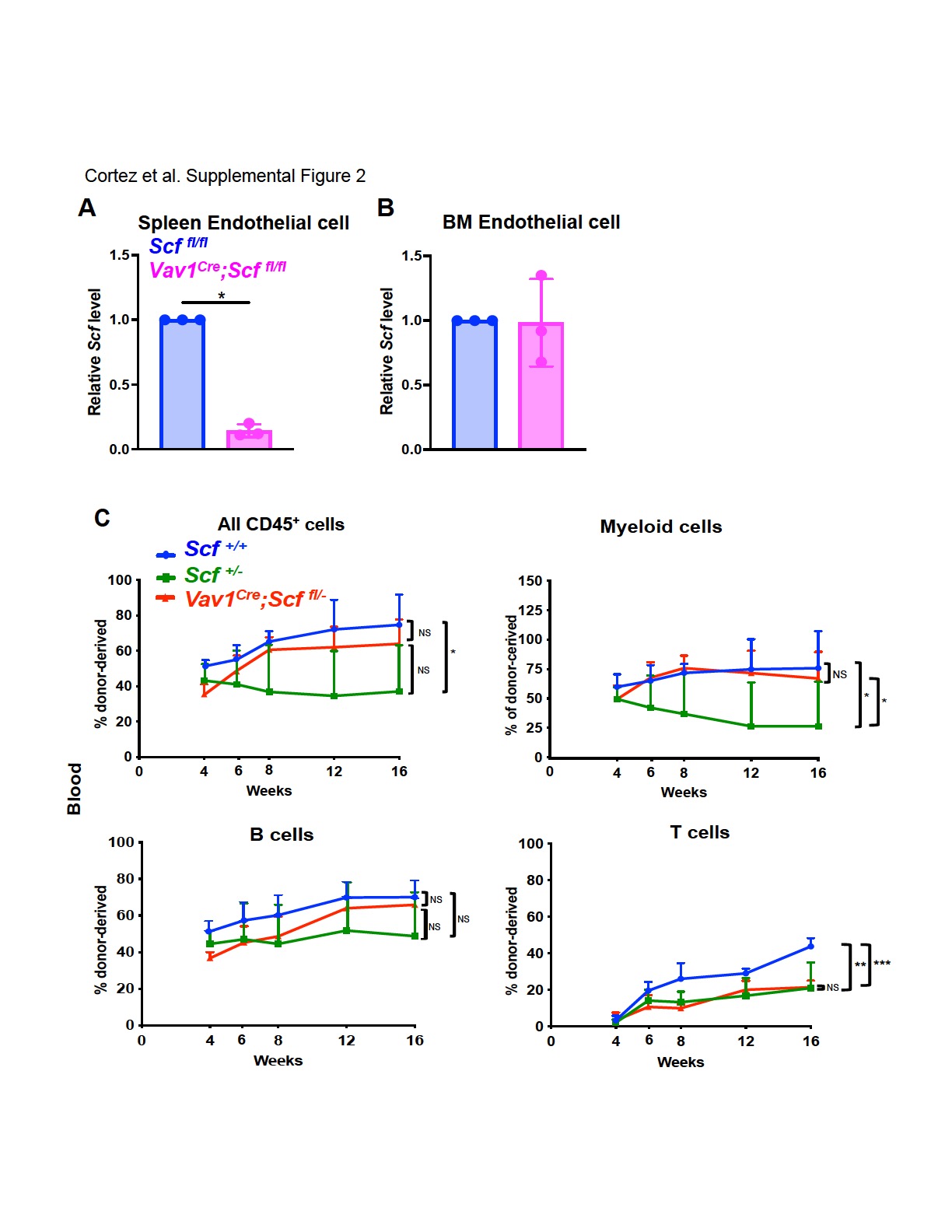

**Supplemental Figure 2. *Vav1-Cre*-mediated *Scf* deletion reduces *Scf* transcript levels in spleen, but not bone marrow, endothelial cells and does not significantly affect the multilineage reconstituting potential of bone marrow cells upon transplantation into irradiated mice.** (A, B) Relative *Scf* transcript levels in spleen (A) and bone marrow (B) endothelial cells from *Vav1^Cre^; Scf ^fl/fl^* and *Scf ^fl/fl^* control mice (a total of 3 mice per genotype from 3 independent experiments; each dot represents a different mouse). (C) Donor-derived reconstitution of CD45^+^ cells, myeloid, B, and T cells in the blood of recipient mice after competitive transplantation of 500,000 donor bone marrow cells from the femurs and tibias of *Vav1^Cre^; Scf^fl/-^* mice, *Scf^+/+^* mice, and *Scf^+/-^* mice, along with 500,000 competing wild-type cells, into irradiated recipients (5 recipient mice per genotype). Statistical significance was assessed using one-sample *t*-tests followed by Holm-Sidak multiple comparisons adjustments (A, B) or nparLD followed by FDR multiple comparisons adjustments (C). All data represent mean ± standard deviation and statistical tests were 2-sided (*, P < 0.05; **, P < 0.01; ***, P < 0.001).

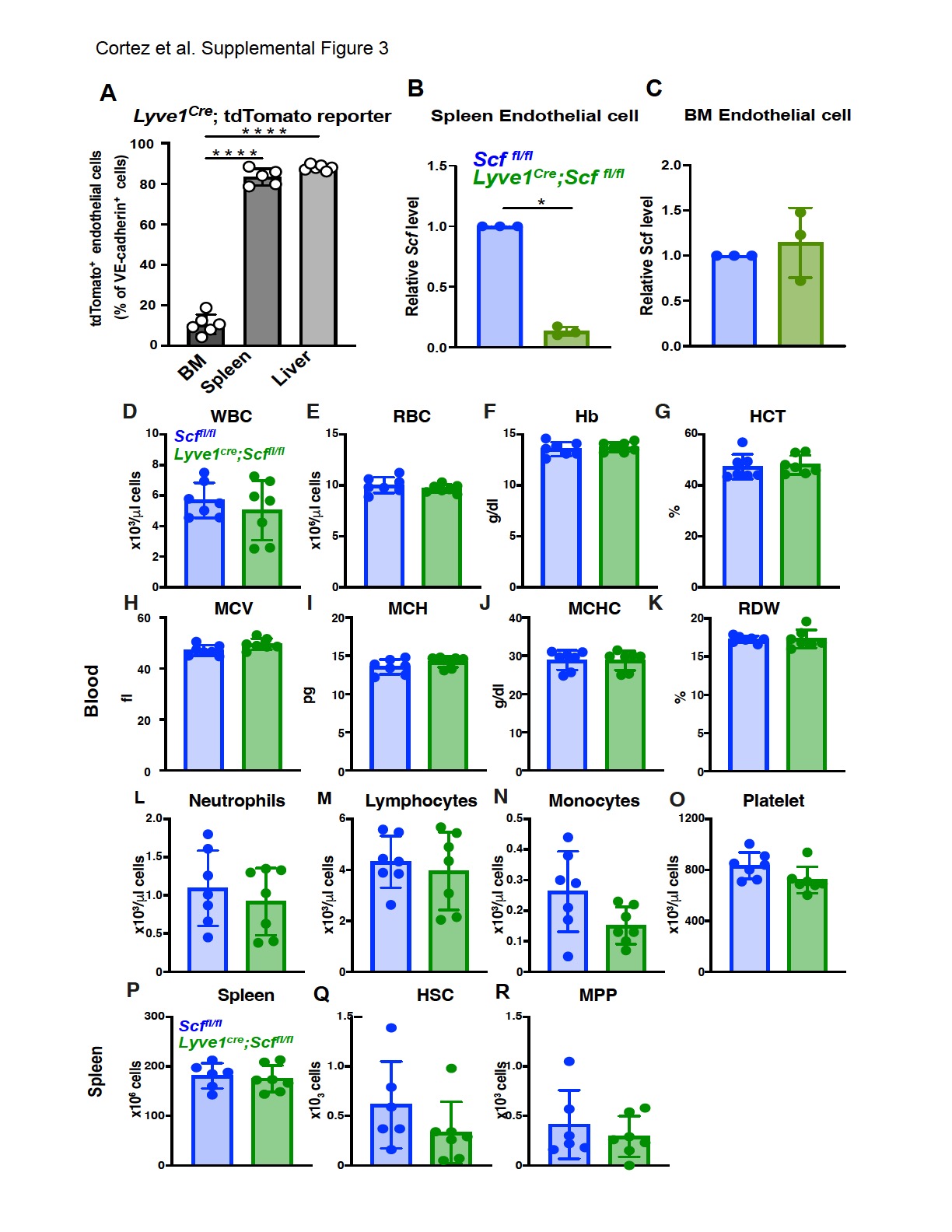

**Supplemental Figure 3.** ***Lyve1-Cre* recombines in spleen and liver, but not bone marrow, endothelial cells and *Lyve1^Cre^; Scf ^fl/fl^* mice have normal steady-state hematopoiesis.** (A) The percentage of VE-cadherin⁺ endothelial cells that were tdTomato+ in bone marrow, spleen, and liver of 3-month-old *Lyve1^Cre^*; *tdTomato* mice (a total of 5–6 mice from four independent experiments). (B, C) Relative *Scf* transcript levels in spleen (B) and bone marrow (C) endothelial cells from *Lyve1^Cre^; Scf ^fl/fl^* and *Scf ^fl/fl^* control mice (a total of 3 mice per genotype from 3 independent experiments). (D-O) White blood cell (D) and red blood cell (E) counts, hemoglobin levels (F) and hematocrit (G), mean corpuscular volume (H), mean corpuscular hemoglobin (I), mean corpuscular hemoglobin concentration (J), red cell distribution width (K), and neutrophil (L), lymphocyte (M), monocyte (N) and platelet (O) counts in the blood of *Lyve1^Cre^; Scf ^fl/fl^* and littermate control (*Scf ^fl/fl^*) mice. (P-R) Spleen cellularity (P) as well as the numbers of HSCs (Q) and MPPs (R) in the spleen of *Lyve1^Cre^; Scf ^fl/fl^* and littermate control mice. Data are from a total of 6-7 mice from 4 independent experiments). All data represent mean ± standard deviation and each dot represents a different mouse. Statistical significance was assessed using a linear mixed-effects model followed by Dunnett’s multiple comparisons adjustment (A) one-sample *t*-tests followed by Holm-Sidak’s multiple comparisons adjustment (B, C) Student’s *t*-tests or Mann-Whitney tests followed by Holm-Sidak’s multiple comparisons adjustments (D-O, and Q-R) or a Student’s *t*-test (P). All statistical tests were 2-sided.

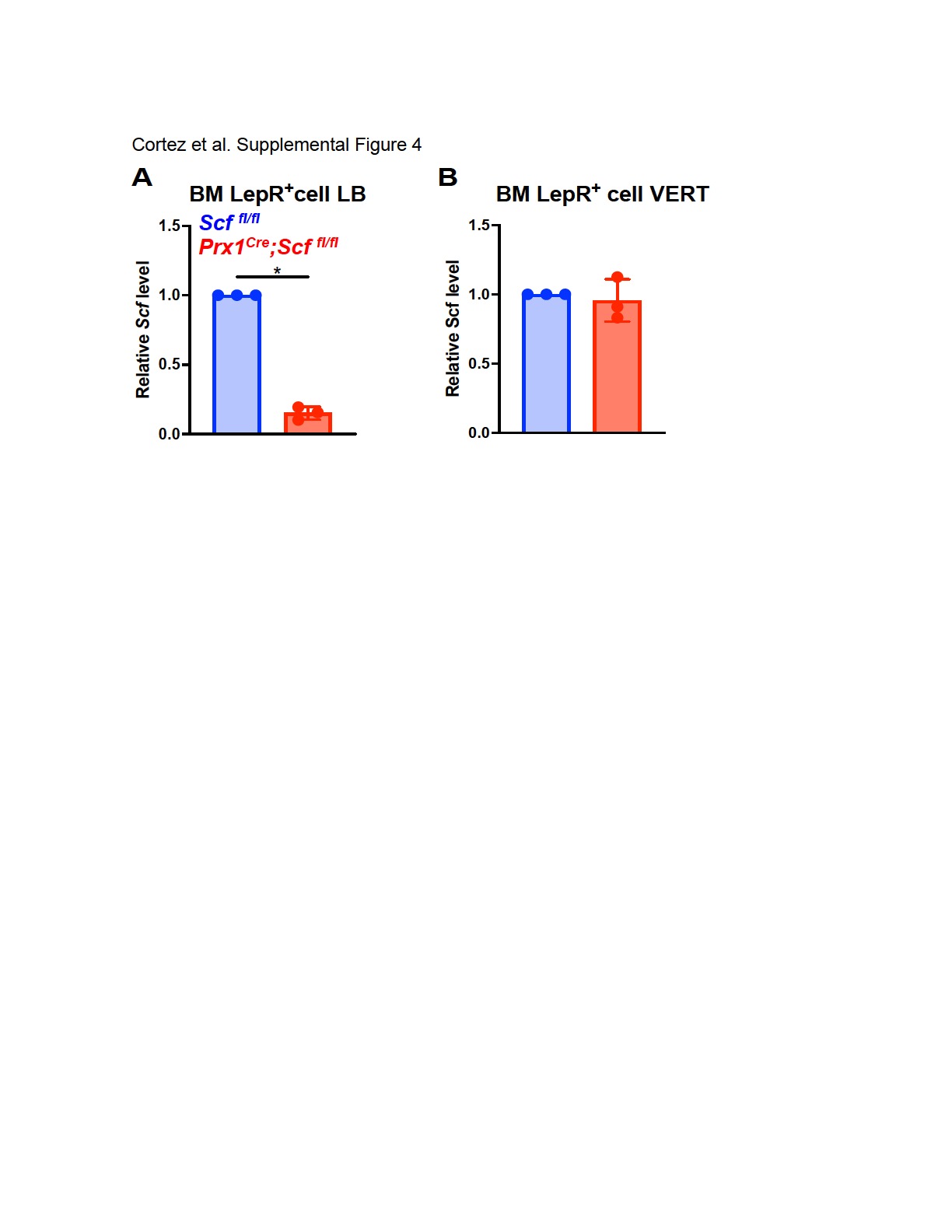

**Supplemental Figure 4. *Prx1-Cre*-mediated recombination of *Scf* reduces *Scf* transcript levels in LepR⁺ bone marrow stromal cells in long bones (femurs and tibia) but not vertebrae.** (A, B) Relative *Scf* transcript levels in LepR⁺ bone marrow stromal cells from long bones (A) versus vertebrae (B) of *Prx1^Cre^; Scf ^fl/fl^* and *Scf ^fl/fl^* control mice (a total of 3 mice per genotype from 3 independent experiments). All data represent mean ± standard deviation and each dot represents a different mouse. Statistical significance was assessed using one-sample *t*-tests followed by Holm-Sidak’s multiple comparisons adjustment (A, B).

**Supplemental Table 1.** Antibodies and viability reagents used for flow cytometry.

| **ANTIBODY OR REAGENT** | **FLUOROCHROME** | **CATALOG NUMBER** |
| --- | --- | --- |
| CD150 (TC15-12F12.2; BioLegend) | BV421 | 115925 |
| CD2 (RM2-5; Tonbo Biosciences) | FITC | 35-0021 |
| CD3 (17A2; BioLegend) | FITC | 100203 |
| CD5 (53-7.3; BioLegend) | FITC | 100605 |
| CD8α (53-6.7; Tonbo Biosciences) | FITC | 35-0081 |
| B220/CD45R (RA3-6B2; Tonbo Biosciences) | FITC | 35-0452 |
| Ter119 (TER-119; Tonbo Biosciences) | FITC | 35-5921 |
| Gr-1/Ly-6G/Ly-6C (RB6-8C5; Tonbo Biosciences) | FITC | 35-5931 |
| CD127/IL-7Rα (A7R34; Tonbo Biosciences) | PE | 50-1271 |
| Viability reagent (BD Pharmingen) | Propidium iodide (PI) | 556463 |
| Sca-1/Ly-6A/E (D7; Thermo Fisher Scientific/eBioscience) | PE-Cyanine7 | 25-5981-82 |
| CD48 (HM48-1; BioLegend) | Alexa Fluor 700 | 103425 |
| CD135/Flt3 (A2F10; BioLegend) | APC | 135310 |
| c-Kit/CD117 (2B8; Thermo Fisher Scientific/eBioscience) | APC-eFluor 780 | 47-1171-82 |
| CD105/Endoglin (MJ7/18; BD Biosciences) | BV421 | 562760 |
| CD34 (RAM34; Thermo Fisher Scientific/eBioscience) | FITC | 11-0341-82 |
| CD16/32/FcγRIII/II (93; BioLegend) | BV510 | 101333 |
| CD2 (RM2-5; Tonbo Biosciences) | PE | 50-0021 |
| CD3 (17A2; Tonbo Biosciences) | PE | 50-0032 |
| CD5 (53-7.3; BioLegend) | PE | 100607 |
| CD8α (53-6.7; Tonbo Biosciences) | PE | 50-0081 |
| B220/CD45R (RA3-6B2; Tonbo Biosciences) | PE | 50-0452 |
| Ter119 (TER-119; Tonbo Biosciences) | PE | 50-5921 |
| Gr-1/Ly-6G/Ly-6C (RB6-8C5; Tonbo Biosciences) | PE | 50-5931 |
| CD41 (MWReg30; BioLegend) | Alexa Fluor 700 | 133926 |
| CD150 (TC15-12F12.2; BioLegend) | APC | 115909 |
| CD71/Transferrin receptor (RI7217; BioLegend) | BV421 | 113813 |
| Gr-1/Ly-6G/Ly-6C (RB6-8C5; BioLegend) | BV510 | 108437 |
| CD43 (S7; BD Biosciences) | PE | 553271 |
| Ter119 (TER-119; BioLegend) | PE-Cyanine7 | 116222 |
| CD3 (17A2; Thermo Fisher Scientific/eBioscience) | Alexa Fluor 700 | 56-0032-82 |
| IgM (II/41; Thermo Fisher Scientific/eBioscience) | APC | 17-5790-82 |
| Mac-1/CD11b (M1/70; Thermo Fisher Scientific/eBioscience) | APC-eFluor 780 | 47-0112-82 |
| LepR (polyclonal; R&D Systems; biotinylated) | Biotin | AF497 |
| Streptavidin (BioLegend) | BV421 | 405226 |
| CD45 (30-F11; Thermo Fisher Scientific/eBioscience) | APC-eFluor 780 | 47-0451-82 |
| Ter119 (TER-119; Thermo Fisher Scientific/eBioscience) | APC-eFluor 780 | 47-5921-82 |
| Ghost Dye Red 780 viability dye (Tonbo Biosciences) | Ghost Dye Red 780 | 13-0865-T100 |
| CD31 (390; BioLegend) | PE/Cyanine7 | 102418 |
